# Transcriptional landscape of soybean (*Glycine max*) embryonic axes during germination in the presence of paclobutrazol, a gibberellin biosynthesis inhibitor

**DOI:** 10.1101/480814

**Authors:** Rajesh K. Gazara, Eduardo A. G. de Oliveira, Antônia Elenir A. Oliveira, Thiago M. Venancio

## Abstract

Gibberellins (GA) are key positive regulators of seed germination. Although the GA effects on seed germination have been studied in a number of species, little is known about the transcriptional reprogramming modulated by GA during this phase in species other than *Arabidopsis thaliana*. Here we report the transcriptome analysis of soybean embryonic axes during germination in the presence of paclobutrazol (PBZ), a GA biosynthesis inhibitor. We found a number of differentially expressed cell wall metabolism genes, supporting their roles in cell expansion during germination. Several genes involved in the biosynthesis and signaling of other phytohormones were also modulated, indicating an intensive hormonal crosstalk at the embryonic axis. We have also found 26 photosynthesis genes that are up-regulated by PBZ at 24 hours of imbibition (HAI) and down-regulated at 36 HAI, which led us to suggest that this is part of a strategy to implement an autotrophic growth program in the absence of GA-driven mobilization of reserves. Finally, 30 transcription factors (mostly from the MYB, bHLH and bZIP families) that are down-regulated by PBZ and are likely downstream GA targets that will drive transcriptional changes during germination.

## INTRODUCTION

Gibberellins (GAs) constitute a large family of diterpenoid compounds that are ubiquitous in higher plants. Some GAs regulate processes such as seed germination, root and stem elongation, leaf expansion, flower and fruit development [1, 2]. Seed germination typically starts with imbibition and ends with testa rupture, followed by emergence of the embryonic axis [3]. During this relatively short period, metabolic activity resumes, mitochondria and DNA damaged during desiccation are repaired, stored mRNAs are translated or degraded and new transcriptional programs are activated. This complex series of interconnected events is fueled by the mobilization of stored reserves and gradually shifts towards photosynthesis and autotrophic growth [4-6].

Over the past decades, seminal studies unequivocally demonstrated the role of GA in promoting seed germination [7], in particular because GA-deficient mutants (e.g. *ga1-3* and *ga2-1*) often require exogenous GA to germinate [8, 9]. Further, the inhibition of radicle emergence in the presence of GA biosynthesis inhibitors (e.g. uniconazole and paclobutrazol, PBZ) indicates that GA is essential for seed germination [10-12]. PBZ is a plant growth retardant that blocks GA biosynthesis by inhibiting kaurene oxidase [13]. Other key GA biosynthesis enzymes are GA20- and GA3-oxidases (GA20ox and GA3ox, respectively), whereas GA2-oxidases (GA2ox) inactivate GA. During late germination, GA is synthesized at the radicle, hypocotyls and micropylar endosperm [14]. GA is recognized by soluble receptors of the GIBBERELLIN INSENSITIVE DWARF1 (GID1) family [15], which comprises the subfamilies GID1ac and GID1b in eudicots. Although very similar at the primary sequence level, different lines of evidence indicate that these subfamilies are functionally divergent [2, 16, 17]. The GA–GID1 complex promotes the degradation of DELLA transcriptional repressors via the 26S proteasome pathway [18]. Further, enhanced germination has been reported in loss-of-function DELLA-mutants [19]. GA is also notorious for its antagonistic interactions with ABA, a well-known seed germination inhibitor. In addition, GA has also been proposed to positively interact with brassinosteroids (BRs) and ethylene, which are ABA antagonists during seed germination [19-21].

During seed germination, GA enhances embryo growth by promoting cell elongation and weakening of the surrounding tissues [14, 19]. Several genes regulated by GA or DELLA have been identified during *Arabidopsis* seed germination, seedling and floral development [14, 22-24]. In addition, various genes related to hormone pathways and cell wall metabolism were modulated by GA [14, 22]. Despite the valuable information accumulated on the biochemical details of GA signaling and interactions with other hormones, little is known about the transcriptional programs driven by GA in germinating seeds of species other than *A. thaliana*. To date, only one report investigated the transcriptome of embryonic axes during soybean (*Glycine max*) germination [25]. Although this study showed a conspicuous activation of GA biosynthesis genes, it does not allow one to distinguish GA-driven transcriptional alterations. In the present work, we report the transcriptome of soybean embryonic axes during seed germination in the presence of the GA biosynthesis inhibitor PBZ, aiming to uncover the genes that are regulated by GA. We show that PBZ: 1) up-regulates several photosynthesis genes; 2) modulates the expression of numerous genes involved in the biosynthesis, signaling and transport of other hormones, suggesting an intensive hormonal cross-talk during germination; 3) modulates the expression of several genes encoding cell wall modifying enzymes, supporting their roles in embryo cell expansion during germination and; 4) represses several transcription factors (TFs) in a time-specific fashion, indicating that these TFs might drive the transcriptional reprogramming mediated by GA during germination.

## RESULTS AND DISCUSSION

### Transcriptome sequencing and functional analysis of differentially expressed genes

We conducted an initial assay to investigate the effects of PBZ on soybean seed germination. As expected, PBZ administration reduced radicle length, fresh weight and dry weight, resulting in a delay in germination (Supplementary figure S1). Embryonic axes at 12, 24 and 36 hours after imbibition (HAI) were carefully separated from the cotyledons and submitted to RNA extraction, library preparation and sequencing on an Illumina HiSeq 2500 instrument (see methods for details). A total of 18 libraries (three biological replicates, with or without PBZ) were sequenced, resulting in a total of 14 to 67 million reads per sample (Supplementary table S1). High-quality reads were mapped to the soybean reference genome (Wm82.a2.v1) and used for downstream analysis. Overall, 97.2% of the reads mapped to the reference genome (Supplementary table S1). In general, we found good correspondence between the biological replicates (Supplementary figure S2) and high pair-wise correlations (0.95 to 0.99) (Supplementary table S2). Genes with RPKM (Reads Per Kilobase per Million mapped reads) greater than or equal to 1 were considered expressed. In total, 29,204, 29,467, 31,065, 30,887, 32,636 and 32,466 genes were found to be expressed in 12C (control), 12P (PBZ), 24C, 24P, 36C and 36P, respectively (Figure 1A). Approximately 62.43% of the soybean protein-coding genes (34,990 genes) were expressed in at least at one time point (Supplementary table S3), which is comparable to a previously published soybean germination transcriptome [25].

**Figure 1.**
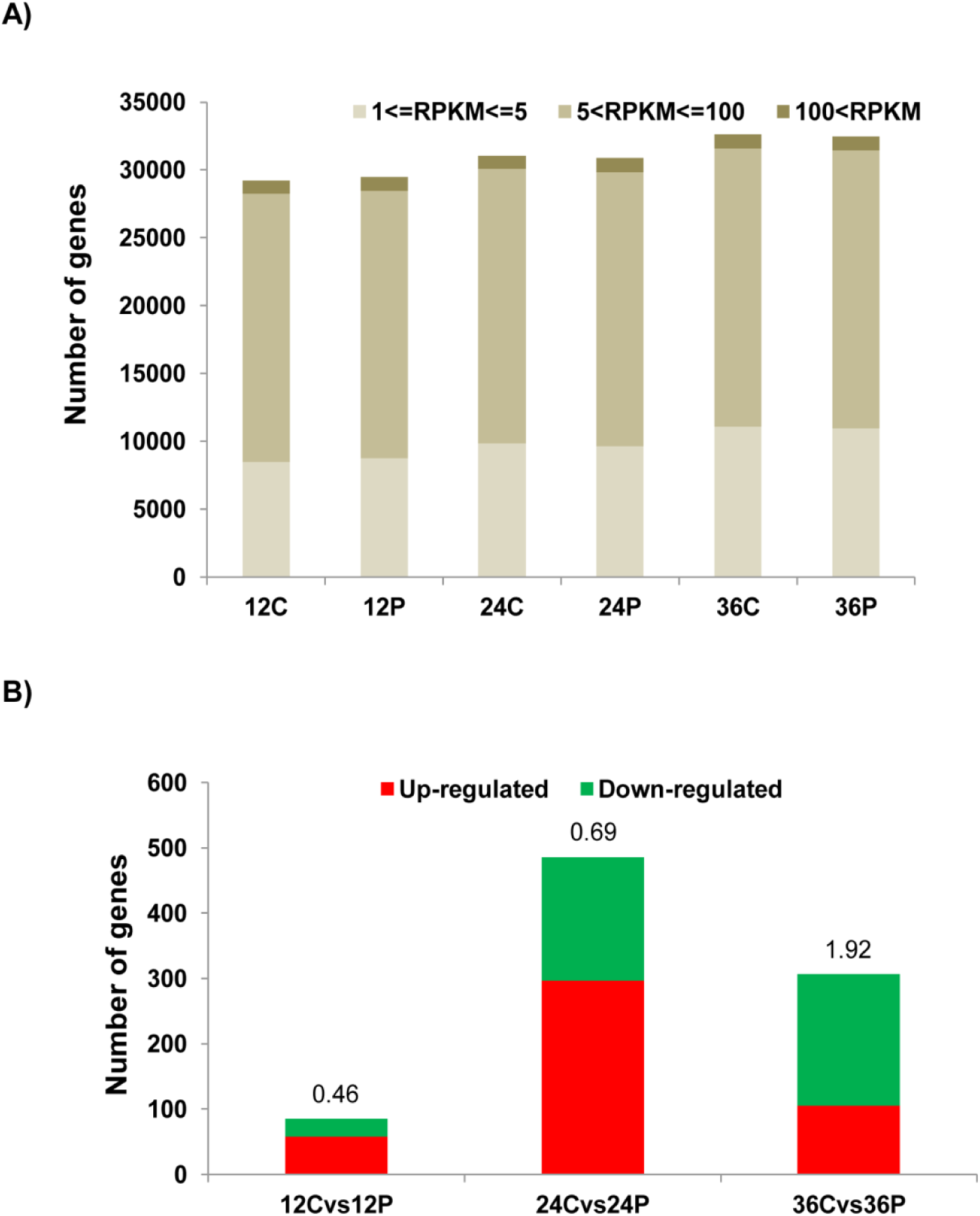
Gene expression profiling during seed germination. **A)** Number of expressed genes (RPKM ≥ 1) and their estimated expression levels in each sample. **B)** Number of DEGs at 12, 24 and 36 HAI. Numbers above the vertical bars stand for the ratio between down- and up-regulated genes. In the x-axes labels, C and P stand for control and PBZ, respectively.

We compared the transcriptional profiles of PBZ-treated seeds at each time point with their respective controls and found a total of 85, 486 and 307 differentially expressed genes (DEGs) at 12, 24 and 36 HAI, respectively (Supplementary table S4). Because PBZ is a GA antagonist, PBZ down- and up-regulated genes (i.e. PBZ-down and PBZ-up, respectively) are likely those induced and repressed by GA. The absolute number of genes down-regulated by PBZ and their ratios to up-regulated genes increased along germination (Figure 1B; Supplementary table S4). In DEG counts, 24 HAI was the most notable time point (297 and 189 up- and down-regulated genes, respectively; Figure 1B). On the other hand, 12 HAI had the lowest number of DEGs (58 and 27 up- and down-regulated genes), indicating that GA transcriptional programs are mostly activated between 12 and 24 HAI and decrease afterwards, when most seeds had completed germination (Supplementary figure S1). Notably, we found 63 genes that are significantly up-regulated at 24 HAI and down-regulated at 36 HAI by PBZ (Supplementary table S5, Supplementary figure S3). About 41% (26 out of 63) of these genes are related with photosynthesis and their up-regulation by PBZ at 24 HAI followed by a down-regulation at 36 HAI might be a strategy to anticipate the transition to autotrophic growth in the absence of energetic resources resulting from proper GA signaling. This gene set encodes chloroplast ATP synthase subunits, RuBisCO, chloroplast ribosomal proteins, DNA-directed RNA polymerase subunit beta (*rpoC1*), YCF3 and several photosystem I and II subunits. Decrease in the expression of plastidial RNA polymerases (i.e. *rpoB* and *rpoC1)* caused aberrant chloroplast development and diminish photoautotrophic growth in *A. thaliana* [26]. *Chloroplast YCF3* encodes a thylakoid protein that is essential for photosystem I complex biogenesis in tobacco [27] and *Chlamydomonas reinhardtii* [28]. The regulation of photosynthesis genes by GA has also been recently demonstrated in rice seedlings under submergence [29]. Interestingly all these 26 genes are nuclear encoded copies of genes that are located in the soybean chloroplast (Reference Sequence: NC_007942). Most of these copies seem to be functional, as they encode proteins with high sequence coverage (68 to 100%) and similarity (78 to 100%) with their plastidial counterparts (Supplementary table S6). Similarly, 17 out of 63 genes encode proteins similar to those encoded by mitochondrial genes (Reference sequence: JX463295) (Supplementary table S6). Collectively, these genes might integrate a system to reduce the dependence on cotyledonary reserves and optimize ATP production. Out of these 43 genes with organellar copies, 41 have been assigned to soybean reference chromosomes, suggesting that they are not annotated as nuclear genes due to contamination of organelle DNA fragments.

### Gene Ontology and KEGG pathway enrichment analysis

Aiming to unravel major trends in the DEG lists, we conducted Gene Ontology (GO) and KEGG pathway enrichment analyses. There was no enrichment of GO terms or KEGG pathways at 12 HAI. In up-regulated genes at 24 HAI, we found a total of 19 enriched GO terms, including terms related with photosynthesis and translation (Supplementary table S7). Three of the GO terms enriched in the genes up-regulated at 24 HAI were also found enriched in the genes down-regulated at 36 HAI, namely “generation of precursor metabolites and energy”, “photosynthesis” and “thylakoid” (Supplementary table S7), providing further support to the results discussed in the previous section.

KEGG pathway enrichment analysis revealed that ‘plant hormone signal transduction’ was enriched in down-regulated genes at 24 HAI and 36 HAI, supporting the regulation of other hormonal pathways by GA, and possibly their cross-talk, during germination (Supplementary table S8). These genes are involved in BR, auxin, jasmonic acid, ABA and cytokinin signaling or biosynthesis. Given their indispensable roles in regulating seed germination, genes related with hormone signaling and biosynthesis are discussed in more detail in the next section. In down-regulated genes at 36 HAI, a number of genes encoding chaperones resulted in the enrichment of the pathway ‘protein processing in endoplasmic reticulum’. Phenylpropanoid biosynthesis genes were enriched in PBZ-down (7 genes) and PBZ-up genes (6 genes) at 24 HAI and 36 HAI, respectively. These genes include β-glucosidases, peroxidases and spermidine hydroxycinnamoyl transferases that might be involved in cell wall modification or oxidative stress response (Supplementary table S8). ‘Biosynthesis of secondary metabolites’ genes were enriched in PBZ-down (15 genes) at 24 HAI and, both in PBZ-up (23 genes) and PBZ-down genes (15 genes) at 36 HAI. Most of these PBZ-up genes encode UDP-glycosyltransferases, cytochrome P450 proteins and brassinosteroid-6-oxidases, whereas PBZ-down genes encode 3-ketoacyl-CoA synthases, 1-amino-cyclopropane-1-carboxylate synthases (ACS) and peroxidases (Supplementary tables S8). ‘Glutathione metabolism’, ‘RNA polymerase’, ‘purine metabolism’, ‘nucleotide excision repair’, ‘pyrimidine metabolism’ and spliceosome pathways were only enriched in up-regulated genes at 24 HAI (Supplementary tables S8). All ‘glutathione metabolism’ DEGs encode glutathione-S-transferases (GSTs) and their up-regulation is related to an increased antioxidant capacity [30]. Increased antioxidant capacity and DNA repair mechanisms at 24 HAI in response to PBZ might be part of a tolerance mechanism to cope with the germination delay, which is in line with a recent study that proposed a link between DNA repair and antioxidant activity in *Medicago truncatula* seed germination and seedling establishment [31].

### Feedback regulation and cross-talk with other hormones

GA biosynthesis can be divided into early (*CPS, KS, KO* and *KAO*) and late (e.g. *GA20ox* and *GA3ox*) stages [1]. While early GA biosynthesis genes are generally not affected by GA [32], a negative GA-mediated feedback mechanism involving the down-regulation of late GA biosynthesis genes and up-regulation of the GA-deactivating *GA2ox* has been proposed as a system to keep balanced GA levels [1]. Although not included by our statistical thresholds, we found *GA3ox* and *GA20ox* genes with greater expression in the presence of PBZ at 24 HAI and 36 HAI (Figure 2, Supplementary table S9), which might indicate a compensating mechanism in response to PBZ. Long- and short-distance GA movement are also critical for developmental processes such as seed germination [33]. Recently, some transporters from the NPF and SWEET families transport GA *in planta* [34, 35]. Curiously, NPF3 transports GA and ABA in *A. thaliana* [35]. We found two *NPF3* genes strongly up-regulated by PBZ at 36 HAI, which is in accordance with the GA-mediated repression of *NPF3* expression [35]. The spatiotemporal expression pattern of NPF3 has been proposed as a key aspect of its functionality [35]. In line with this, recent elegant works in *A. thaliana* showed that GA gradients correlate with cell length in dark-grown hypocotyls [36, 37]. We hypothesize that this might be the case in soybean embryonic axes, particularly in the context of the recently described radicle-derived growth pattern in germinating soybean embryos [38].

**Figure 2.**
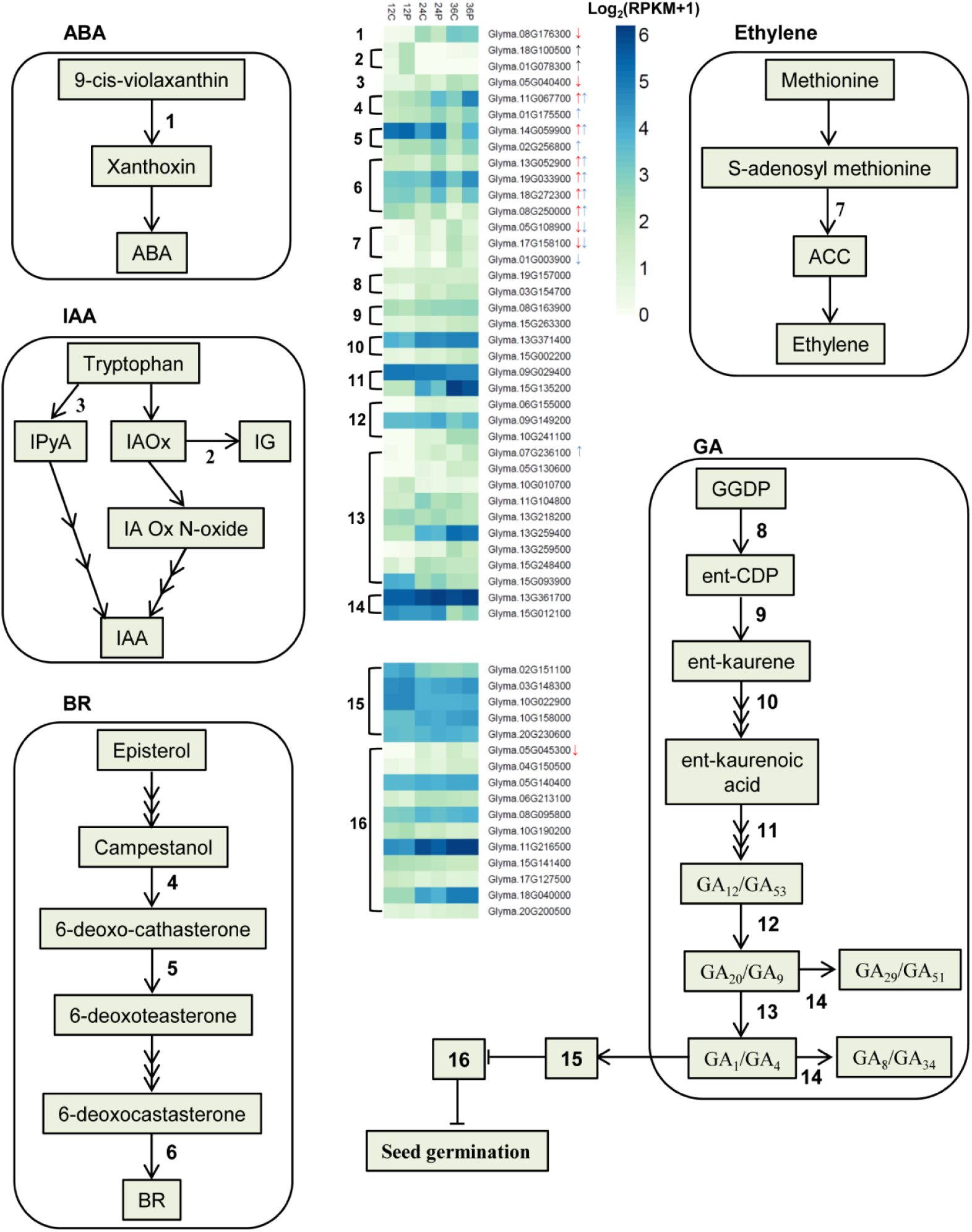
Hormone biosynthesis pathways. Some GA deactivation and signaling genes discussed are also included. Up- and down-regulated genes are shown with up and down arrows. Black, red and blue arrows represent differential expression at 12, 24 and 36 HAI, respectively. Genes without arrows are expressed in at least one condition, although not included by our statistical thresholds. Genes are numbered as follows: 1) nine-cis-epoxycarotenoid dioxygenase 3 (*NCED3*); 2) *SUR2*; 3) tryptophan aminotransferase related 2 (*TAR2*); 4) DWARF4 (*DWF4*); 5) DWARF3 (*DWF3*); 6) brassinosteroid-6-oxidase 2 (*BR6ox2*); 7) 1-amino-cyclopropane-1-carboxylate synthase (*ACS*); 8) ent-copalyl diphosphate synthase (*CPS*); 9) ent-kaurene synthase (*KS*); 10) ent-kaurene oxidase (*KO*); 11) ent-kaurenoic acid oxidase (*KAO*); 12) GA 20-oxidase (*GA20ox*); 13) GA 3-oxidase (*GA3ox*); 14) GA 2-oxidase (*GA2ox*); 15) GIBBERELLIN INSENSITIVE DWARF1 (*GID1*) [*Glyma.02G151100* (GID1b1), *Glyma.10G022900* (GID1b2), *Glyma.03G148300* (GID1b3), *Glyma.10G158000* (GID1c1) and *Glyma.20G230600* (GID1c2); 16) DELLA. Abbreviations: Abscisic Acid (ABA), indole-3-pyruvic acid (IPyA), Indol-3-acetaldoxime (IAOx), Indol-3-acetaldoxime N-oxide (IA Ox N-oxide), indole glucosinolates (IG), Indole-3-acetic acid (IAA), Brassinosteroid (BR), 1-aminocyclopropane-1-carboxylic acid (ACC), geranyl geranyl diphosphate (GGDP), ent-copalyl diphosphate (ent-CDP).

In addition to biosynthesis and transport, we have also investigated GA signaling genes. We found 11 DELLA genes (one PBZ-down at 24 HAI) and all 5 GID1s [17] expressed in at least one time point (Figure 2, Supplementary table S9). Almost all DELLAs showed greater expression in the absence of PBZ (Figure 2, Supplementary table S9). The expression levels of GID1b1, GID1b2 and GID1b3 were greater in PBZ than in controls (except GID1b1 and GID1b3 at 24 HAI), supporting that GID1b is particularly important under low GA concentrations, as previously hypothesized by us and others [2, 17]. Collectively, our results support that the low GA production resulting from PBZ administration activates an intricate system involving GA biosynthesis, signaling and transport genes, probably to minimize the effects of impaired GA production to allow germination to occur.

### Other phytohormones

ABA is the most notorious GA antagonist for its inhibitory effect on seed germination [1, 19]. The regulatory step in ABA biosynthesis is catalyzed by 9-cis-epoxycarotenoid dioxygenase (NCED), which is transcriptionally regulated by positive and negative feedback loops in different species [39, 40]. The ABA receptor (PYL) inhibits the protein phosphatase 2C (PP2C) in the presence of ABA [41]. We found one *NCED3* (*Glyma.08G176300*) and two *PP2Cs* as PBZ-down and one *PYL5* as PBZ-up (Figures 2 and 3, Supplementary table S9). In addition, two ABA transporters, *ABCG40* (up-regulated, *Glyma.19G169400*) and *NRT1.2* (down-regulated, *Glyma.08G296000*) were also differentially expressed upon PBZ treatment (Supplementary table S9). Collectively, these results show that GA modulate different genes involved in ABA biosynthesis, signaling and transport, which might directly interfere with a gradient of GA:ABA ratios along germinating soybean embryonic axes. This GA:ABA dynamics might be involved in the differential cell expansion patterns observed in germinating soybean embryos [38].

**Figure 3.**
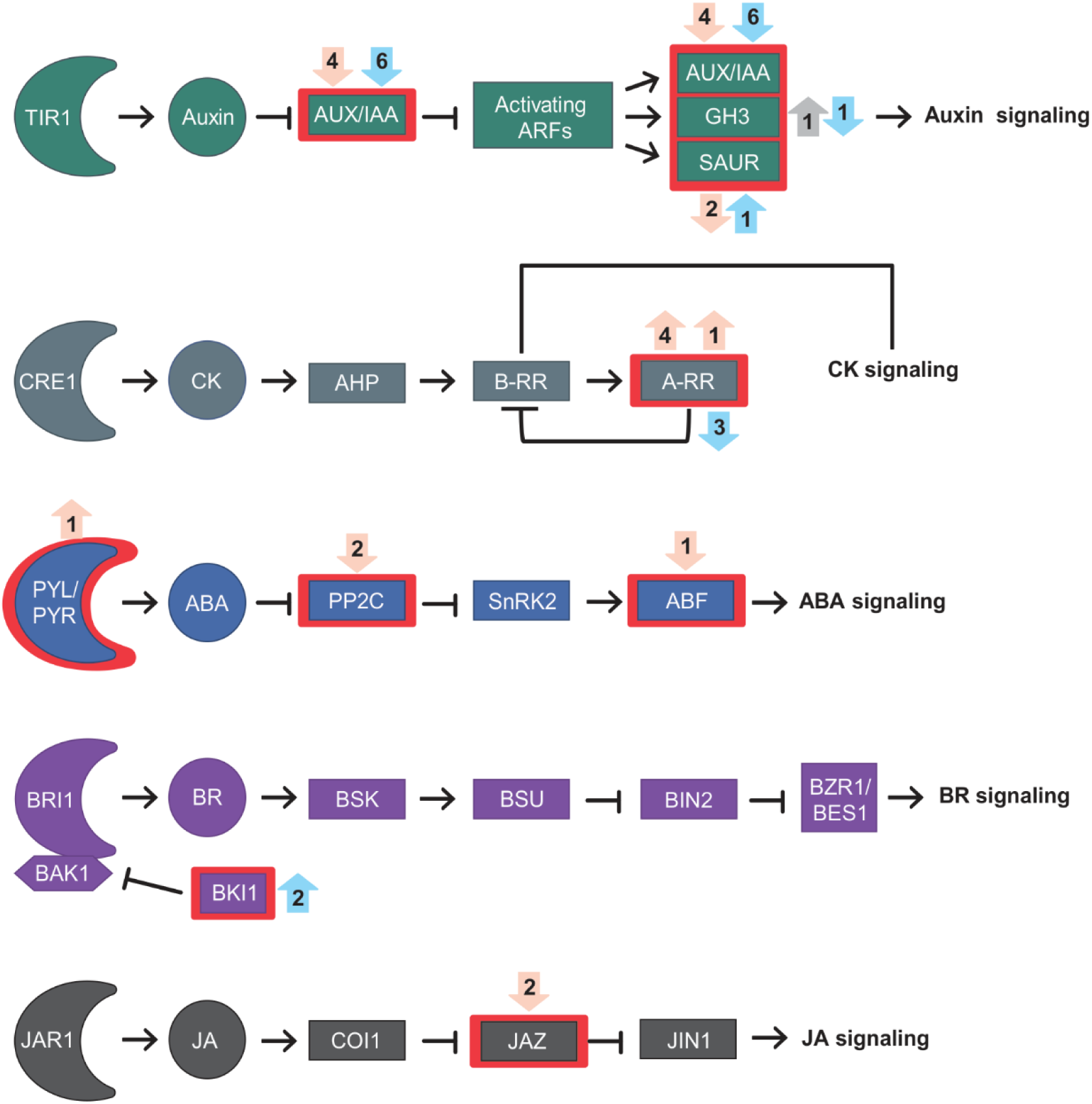
Hormone signal transduction. Rectangles with red lines represent gene families with at least one DEG. Up and down arrows represent PBZ up- and down-regulated genes. Number of DEGs are shown in circles adjacent to the red rectangles. Grey, light orange and light blue arrows represent DEGs at 12, 24 and 36 HAI, respectively. Abbreviations: transport inhibitor response 1 (TIR1); Auxin/Indole-3-Acetic Acid (Aux/IAA); auxin-responsive Gretchen Hagen3 (GH3); small auxin upregulated RNA (SAUR); CYTOKININ RESPONSE 1 (CRE 1); Cytokinin (CK); His-containing phosphotransfer protein (AHP) ;Type-B response regulator (B-RR); Type-A response regulator (A-RR); Pyrabactin Resistance (PYR); PYR-like (PYL); Abscisic acid (ABA); Protein Phosphatase 2C (PP2C); Sucrose non-fermenting 1-related protein kinases subfamily 2 (SnRK2s); Abscisic acid responsive element-binding factor (ABF); Brassinosteroid-insensitive 1 (BRI1); BRI1-associated receptor kinase 1 (BAK1); Brassinosteroid (BR); BRI1 kinase inhibitor (BKI1); Brassinosteroid signaling kinases (BSK); BRI1-suppressor (BSU); brassinosteroid-insensitive 2 (BIN2); Brassinazole-resistant 1 (BZR1); BRI1-ethyl methanesulfonate-suppressor 1 (BES1); JASMONATE RESISTANT1 (JAR1); Jasmonic acid (JA); Coronatine Insensitive1 (COI1); JASMONATE ZIM DOMAIN (JAZ); JASMONATE INSENSITIVE 1 (JIN1).

**Figure 4.**
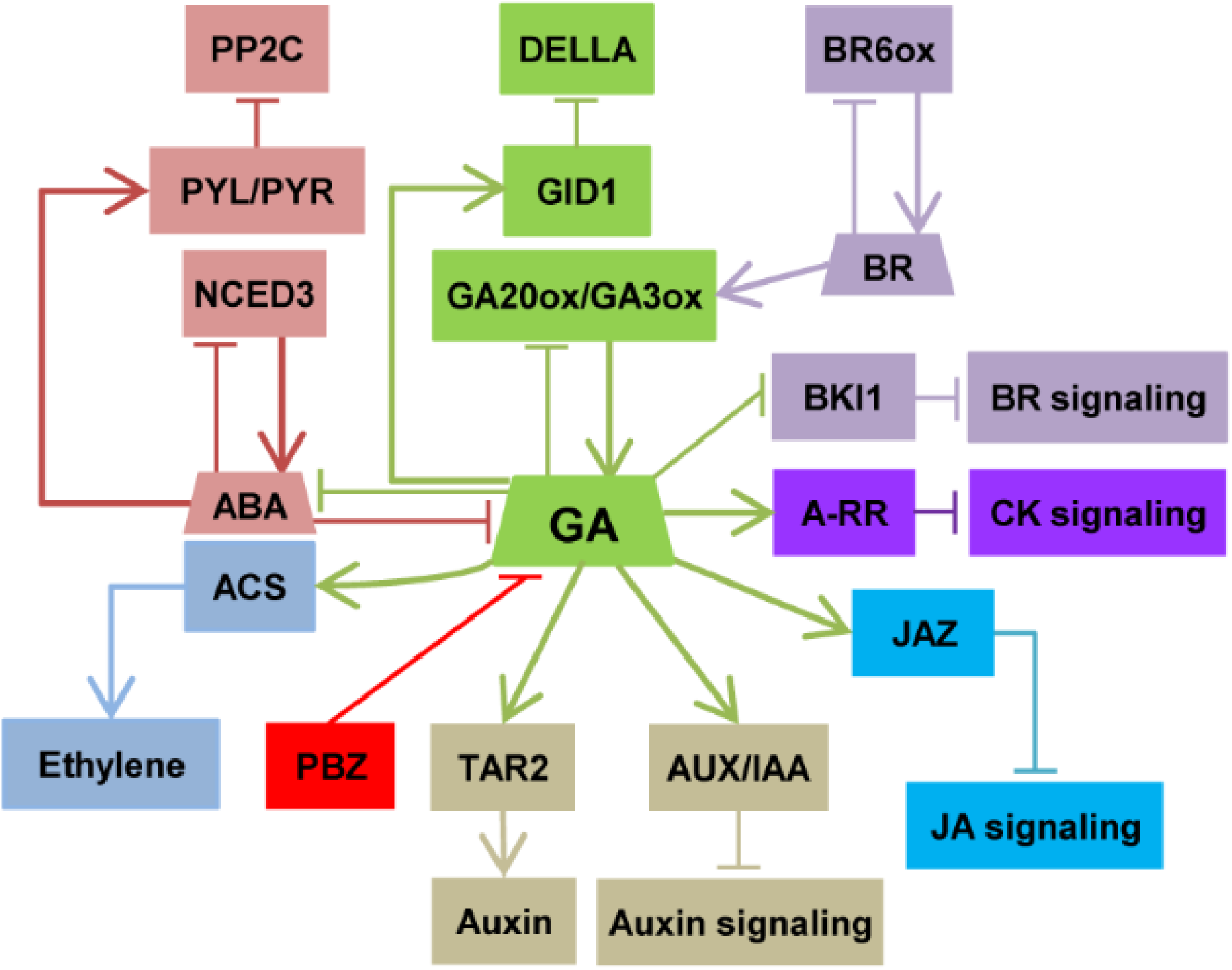
Schematic model of hormonal crosstalk with gibberellin during *G. max* seed germination. The model was derived from a careful literature curation based on differentially expressed genes discussed along the manuscript. Positive interactions are indicated by arrows and T bars indicate repression. Abbreviations: Pyrabactin Resistance (PYR); PYR-like (PYL); Protein Phosphatase 2C (PP2C); Nine-cis-epoxycarotenoid dioxygenase 3 (NCED3); Abscisic acid (ABA); Aminocyclopropane-1-carboxylic acid synthase (ACS); Paclobutrazol (PBZ); Gibberellin (GA); GIBBERELLIN INSENSITIVE DRAWF 1 (GID1); GA 20-oxidase (GA20ox); GA 3-oxidase (GA3ox); Tryptophan aminotransferases 2 (TAR2); Auxin/Indole-3-Acetic Acid (AUX/IAA); Brassinosteroid (BR); BR 6-oxidase (BR6ox); BRI1 kinase inhibitor (BKI1); Type-A response regulator (A-RR); JASMONATE ZIM DOMAIN (JAZ); Jasmonic acid (JA).

GA and ethylene positively interact with each other, promoting seed germination in several species [42]. Multiple lines of evidence, including PBZ administration, support the positive regulation of ethylene biosynthesis and signaling by GA [14, 43-46]. Further, several ethylene biosynthesis genes are expressed in soybean embryonic axes during germination [25]. Accordingly, we found three PBZ-down 1-amino-cyclopropane-1-carboxylate synthase (ACS) genes (Figure 2, Supplementary table S9). ACS catalyzes the first committed and rate-limiting step in ethylene biosynthesis [47]. Our results suggest that up-regulation of ACS by GA is likely a key part of the synergy between GA and ethylene during soybean germination.

Several studies have shown that auxin inhibits or delays seed germination in wheat [48], *Arabidopsis* [49] and soybean [50]. On the other hand, exogenous GA_4_ up-regulated auxin biosynthesis and carrier genes in germinating *Arabidopsis* seeds [14], supporting a complex GA-auxin cross-talk during soybean germination. There are multiple tryptophan-dependent IAA biosynthesis pathways in plants [51]. The tryptophan aminotransferases *TAR1* and *TAR2* convert trp to indole-3-pyruvate (IPA), which is converted to indole acetic acid (IAA) by the YUCCA flavin monooxygenase [52]. Further, *superroot2* (*SUR2*) encodes the cytochrome P450 monooxygenase CYP83B1, involved in glucosinolate biosynthesis and auxin homeostasis [53, 54]. We found two PBZ-up *SUR2* at 12 HAI and one PBZ-down *TAR2* at 24 HAI, indicating that GA promotes IAA production at these time points. We also found one auxin transporter (*PIN*; PBZ-up) and eleven auxin-responsive genes, including seven PBZ-down Auxin/Indole-3-Acetic Acid (Aux/IAA) repressors, small auxin upregulated RNA (SAUR), and the auxin-responsive Gretchen Hagen3 (GH3) family were differentially expressed at least at one of the time-point (Figures 2 and 3, Supplementary table S9). Although apparently conflicting with the promotion of IAA biosynthesis at 12 and 24 HAI, the down-regulation of several *AUX/IAA* genes by PBZ at 24 HAI and 36 HAI suggests that GA represses auxin signaling during late germination. Accordingly, three AUX/IAA genes have been recently demonstrated to promote hypocotyl elongation in *A. thaliana* [55].

BRs typically induce seed germination and BR biosynthesis genes (*DET2, DWF4, DWF3, BR6ox1*, and *ROT3*) are up-regulated when endogenous BR concentrations are reduced [56]. Interestingly, six and eight BR biosynthesis genes were PBZ-up at 24 and 36 HAI, respectively (Figure 2, Supplementary table S9). BR promotes GA biosynthesis by regulating *GA20ox1* and *GA3ox1* expression in *A. thaliana* [57]. Further, GA partially rescued hypocotyl elongation defects resulting from BR deficiency [57]. Our group has proposed that BR signaling regulates cell expansion during soybean germination [25]. Taken together, the up-regulation of BR biosynthesis upon PBZ treatment might be involved in the activation of late GA biosynthesis genes to counter PBZ effects on GA production. This hypothesis also fits the observation that PBZ delays germination without a clear effect on germination rates (Supplementary Figure S1E). Finally, since BR also promotes GA biosynthesis in rice [58], the emergence of this regulatory module probably predates the diversification of monocotyledonous and dicotyledonous species.

Antagonistic interactions between GA and cytokinin (CK) have been reported in different plants [59-61]. Type-A response regulators negatively regulate CK signaling by competing with type-B response regulators for phosphoryl transfer from the upstream *Arabidopsis* Hpt proteins or by interacting with other pathway components [62]. We found four and three PBZ-down type-A response regulators at 24 HAI and 36 HAI, respectively (Figure 3, Supplementary table S9). Since CK biosynthesis genes were not differentially expressed, our results indicate GA antagonizes CK by the up-regulation of negative CK signaling regulators during soybean germination.

In the canonical Jasmonic Acid (JA) signaling pathway, the receptor CORONATINE INSENSITIVE 1 (COI1) interacts with JA and promotes the proteasomal degradation of JASMONATE ZIM-domain (JAZ) repressors [63]. JAZ represses the transcription of JA-responsive genes through interaction with the MYC2 TF and other regulatory proteins [63, 64]. JA and GA perform antagonistic roles in regulating hypocotyl elongation via physical interactions between JAZ and DELLA repressors. In summary, JA-mediated JAZ degradation releases DELLA to repress GA signaling (and hypocotyl elongation), whereas GA-mediated DELLA degradation releases JAZ to inhibit JA responses [64, 65]. We found two PBZ-down *JAZ* genes at 24 HAI (Supplementary table S9), indicating that GA represses JA signaling during germination. Interestingly, *JAZ* up-regulation might constitute an additional layer of JA repression, as GA-promoted DELLA degradation would release JAZ proteins to repress JA signaling, as discussed above.

### Gibberellins regulate cell wall remodeling enzymes

Several genes encoding cell elongation and cell wall remodeling enzymes such as xyloglucan endotransglycosylase/hydrolases (XTH), pectin methylesterases (PME), expansins, pectin lyases, aquaporin and others are induced by GA in *Arabidopsis* and tomato seed germination [14, 22, 66-68]. We found a number of these cell wall remodeling genes as differentially expressed (Figure 5A). Peroxidases and glycosyl hydrolases (GHs) also play active role in cell wall loosening [69, 70]. Accordingly, nine and eight peroxidases and GHs were differentially expressed, respectively. Genes involved in pectin metabolism were also modulated by PBZ (Figure 5A), suggesting that this process is also under GA regulation during germination. We also found other cell wall related DEGs, such as arabinogalactan-proteins, fasciclin-like AGPs, hydroxyproline (Hyp)-rich glycoproteins, and proline- or glycine-rich proteins, which play important roles in cell proliferation [71-73] and expansion [74]. Several of those genes are also GA-responsive in cucumber, maize and barley [75-77]. Importantly, 30 out of 44 cell wall DEGs were PBZ-down, supporting that the notorious effect of GA in promoting cell elongation.

**Figure 5.**
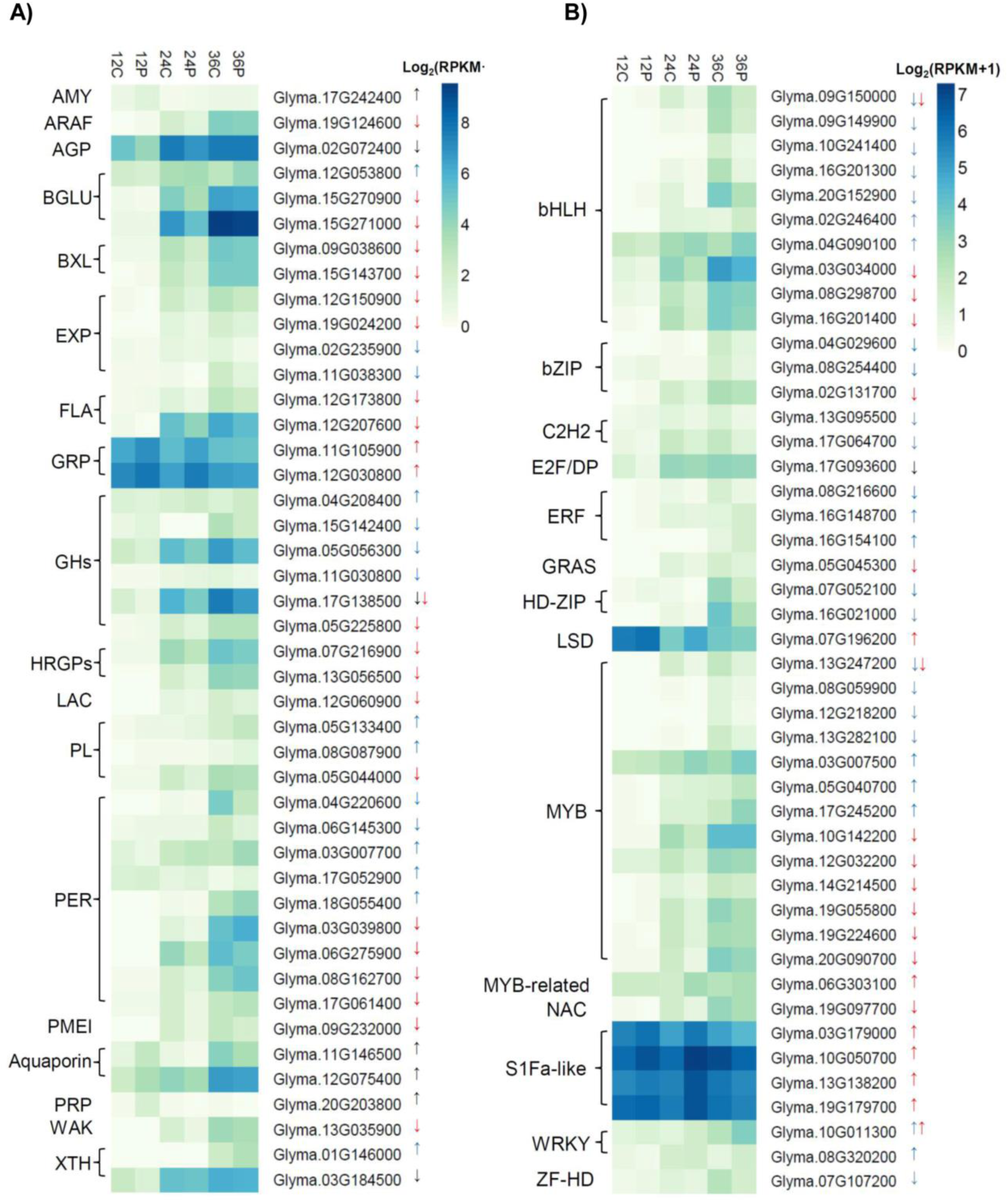
Genes encoding differentially expressed cell-wall remodeling enzymes. (**A) and transcription factors** (**B).** Up- and down-regulated genes are shown with up and down arrows. Black, red and blue arrows represent DEGs at 12, 24 and 36 HAI, respectively. Abbreviations: alpha amylase-like (AMY); alpha-L-arabinofuranosidase (ARAF); arabinogalactan protein (AGP); beta glucosidase (BGLU); beta-xylosidase (BXL); expansin (EXP); FASCICLIN-like arabinogalactan-protein (FLA); glycine-rich protein (GRP); Glycosyl hydrolase family protein (GH); hydroxyproline-rich glycoprotein family protein (HRGP); laccase (LAC); Pectin lyase-like superfamily protein (PL); Peroxidase superfamily protein (PER); pectin methylesterase inhibitor superfamily protein (PMEI); proline-rich protein (PRP); wall associated kinase (WAK); xyloglucan endotransglucosylase/hydrolase (XTH); basic helix-loop-helix (bHLH); Basic Leucine Zipper (bZIP) ; C2H2 zinc finger (C2H2); Ethylene response factor (ERF); GRAS (gibberellin insensitive (GAI), Repressor of ga1-3 (RGA), SCARECROW-LIKE 3 (SCR) gene family; Homeodomain-leucine zipper (HD-ZIP); LESION SIMULATING DISEASE (LSD); Myelobastosis (MYB); Zinc finger Homeodomain (ZF-HD); No apical meristem (NAM), ATAF, and CUC (cup-shaped cotyledon) (NAC) family.

### Transcription factor genes modulated by paclobutrazol are likely drivers of GA-mediated transcriptional reprogramming

Because seed germination is mainly regulated by the embryonic axis, we have specifically investigated the differential expression of TFs in this tissue, as they might be major drivers of the GA transcriptional programs. A total of 45 TFs were differentially expressed upon PBZ treatment. Strikingly, one, 18 and 23 TFs were differentially expressed exclusively at 12, 24 and 36 HAI, respectively (Figure 5B, Supplementary Table S3). This pattern indicates that differentially expressed TFs play specific roles at different germination times. Further, most of the differentially expressed TFs (66.7%) were down-regulated by PBZ and likely comprise regulators that are downstream of GA (Figure 5B, Supplementary Table S3). The TF families with the greatest number of down-regulated members were MyB (myelobastosis; 10 down), bHLH (basic helix-loop-helix; 8 down) and bZIP (basic leucine zipper domain; 3 down), which is in line with previous studies in soybean [25] and *A. thaliana* [22], which showed that MyB and bHLH are among the mostly activated TF families during germination. Interestingly, five and six of the PBZ-down MYB and bHLH genes, respectively, were also differentially expressed in a time-dependent manner during soybean germination [25], further supporting that GA coordinate the transcription of specific TFs at different HAI. Conversely, S1Fa-like (4 up) and WRKY (3 up) were the families that were most represented among PBZ-up TFs (Figure 5B, Supplementary Table S3). S1Fa-like is a poorly-studied TF family that has been associated with photomorphogenesis [78]. Remarkably, all four soybean S1Fa-like TFs were strongly up-regulated by PBZ at 24 HAI, indicating that they might be part of the regulatory system to activate photosynthetic growth in response to low GA concentrations, as discussed above. Photomorphogenesis is regulated by a complex pathway involving GA and light in *A. thaliana* seedlings [79, 80]. Nevertheless, no PIF or HY5 genes, which encode important regulators of photomorphogenesis, were modulated by PBZ.

### Comparison with *A. thaliana* GA-responsive genes

Ogawa *et al* identified a total of 230 and 127 up- and down-regulated genes during germination of *A. thaliana ga1-3* seeds upon GA treatment [14]. Other study, also in *A. thaliana*, reported DEGs in imbibed seeds and developing flowers of wild type, *ga1-3*, and a quintuple DELLA null mutant (*ga1 rga gai rgl1 rgl2*) [22]. This latter study identified 541 and 571 up- and down-regulated GA-responsive genes in imbibed seeds. It is important to mention that Ogawa et al. used a microarray platform representing ∼8,200 genes, while Cao et al. used one covering ∼23,000 genes. This difference is likely an important factor accounting for the differences in DEG numbers between these studies. Overall, these studies have an overlap of 109 GA-up genes and 90 GA-down genes. Importantly, a significant fraction of these genes are also regulated by DELLA [22].

Although *A. thaliana* and soybean are distantly related and their seeds are remarkably different, we investigated the conservation of the DEGs identified in *A. thaliana* described above with the ones reported here using BLASTP (minimum query coverage and similarity of 50%). We found 178 and 124 differentially expressed soybean orthologs for 122 and 84 *A. thaliana* GA-up and GA-down genes, respectively. These soybean gene sets were named GA-up-orthologs and GA-down-orthologs, respectively. Curiously, a significant part (47.19% and 55.66% of the GA-up-orthologs and GA-down-orthologs, respectively) of these genes are modulated in opposite directions in the two species (Supplementary Table S10). Nevertheless, most of the genes related with cell-wall modification, GSTs, auxin responsive genes (AUX/IAA and SAUR), oxidoreductases (aldo-ketoreductases), and transferases are modulated in same directions in soybean and *A. thaliana*, whereas genes modulated in opposite directions between the species encode HSPs, cytochrome p450, serine carboxypeptides, late embryogenesis proteins and flavonol synthase/flavanone 3-hydroxylase (Supplementary Table S10). Proportionally and in absolute numbers, 24 HAI is the stage with the most conserved DEG profile between the two species. Further, 351 out of the 468 soybean DEGs without a DEG ortholog in *A. thaliana* do have orthologs in the *A. thaliana* genome, indicating that a several orthologous genes are differentially regulated in the two species. Finally, in addition to the evolutionary distance, there are also important technical aspects that require consideration. The *A. thaliana* studies used microarrays to investigate modulated genes in *ga1-3* mutants either upon treatment with exogenous GA [14] or in contrast with wild type seeds during germination [22]. Here we analyzed an RNA-Seq transcriptome of embryonic axes of germinating soybean seeds treated with PBZ. Both experimental designs have limitations; even the *A. thaliana ga1-3* dry seeds have bioactive GA from the GA treatment used to rescue parental fertility of mutant plants [14]. In addition, administration of exogenous GA may have unintended effects due to locations and concentrations different from those found under natural conditions. On the other hand, while allowing the investigation with more natural GA concentrations and locations, chemical inhibition of GA biosynthesis probably does not shutdown GA signaling completely. Further, it is not unreasonable to expect that the inhibitor effects might be overcome after some time, for example by an increase in the levels of GA biosynthesis enzymes. A more detailed picture of the interspecies conservation of GA-driven transcriptional programs will be clearer when more species are studied using state-of-the-art RNA-Seq technologies.

## MATERIAL AND METHODS

### Plant material and growth conditions

*G. max* seeds (BRS-284, from EMBRAPA, Brazil) were used in this study. Seeds were surface sterilized with 70% ethanol for 1 minute and with commercial bleach (1% v/v) for 3 minutes, followed by three washes with sterile distilled water (30 seconds per wash). Seeds were germinated in 15 cm Petri dishes with 2 g of sterile cotton in two conditions: in the presence of 30 ml of sterile water (control) or sterile water with 200 µM paclobutrazol (Sigma Aldrich). Seeds were allowed to germinate in an incubation chamber at 28°C and 12/12h photoperiod (dark/light). We used three plates per sample, with 20 seeds per plate. Embryonic axes from dry seeds were also collected. For total RNA extraction, seeds were harvested at 12, 24 and 36 HAI in control and PBZ treated conditions. Embryonic axes were separated from cotyledons and immediately placed in RNA*later*^™^ (Qiagen) until RNA extraction. RNA was extracted from harvested embryonic axes using RNeasy Plant Mini Kit (Qiagen) according to manufacturer instructions. Three independent biological replicates of each condition were used.

### RNA purification, sequencing and analysis

RNA-Seq libraries were prepared using the TruSeq RNA Sample Preparation Kit v2 and submitted to 1×100bp single-end sequencing on a HiSeq 2500 instrument at LaCTAD (UNICAMP, Campinas, Brazil). Read quality was assessed by FastQC (http://www.bioinformatics.babraham.ac.uk/projects/fastqc/). Reads were aligned on *G. max* cv. Williams 82 reference genome version 2 (Wm82.a2.v1) using novoalign (V3.06.05; http://www.novocraft.com). Gene expression levels were calculated with cufflinks v2.1.1 [81] and normalized by reads per kilobase of transcript per million mapped reads (RPKM). Genes with RPKM greater than or equal to one were considered expressed. The differential expression between Control vs PBZ at 12 HAI, 24 HAI and 36 HAI were determined by cuffdiff v2.2.1 [81]. Genes with at least two-fold difference in expression and q-value ≤ 0.05 were considered differentially expressed. Enrichment of Gene Ontology (GO) term was performed using agriGO (v2.0) with hypergeometric test, corrected by the Hochberg FDR method (FDR ≤ 0.05) [82]. Redundant GO terms were removed with REViGO [83]. KOBAS 3.0 [84] was used to assess the enrichment of DEGs in KEGG pathways (Fisher’s exact test, *P* < 0.05). The list of expressed genes (i.e. RPKM ≥ 1) were used as the background set for GO and KEGG enrichment analyses. *G. max* TFs were obtained from the Plant Transcription Factor Database (PlantTFDB) [85]. The datasets generated in this study have been deposited in the NCBI Gene Expression Omnibus database, under the accession number GSE112872.

## Supporting information

## ACKNOWLEDGEMENTS

This work was supported by Fundação Carlos Chagas Filho de Amparo à Pesquisa do Estado do Rio de Janeiro (FAPERJ; grants E-26/010.002019/2014 and E-26/102.259/2013), Coordenação de Aperfeiçoamento de Pessoal de Nível Superior - Brasil (CAPES; Finance Code 001) and Conselho Nacional de Desenvolvimento Científico e Tecnológico (CNPq). We thank the Life Sciences Core Facility (LaCTAD) of State University of Campinas (UNICAMP) for library preparation and RNA sequencing. The funding agencies had no role in the design of the study and collection, analysis, and interpretation of data and in writing.

