## Supplementary material for "Transcriptional landscape of soybean (*Glycine max*) embryonic axes during germination in the presence of paclobutrazol, a gibberellin biosynthesis inhibitor"

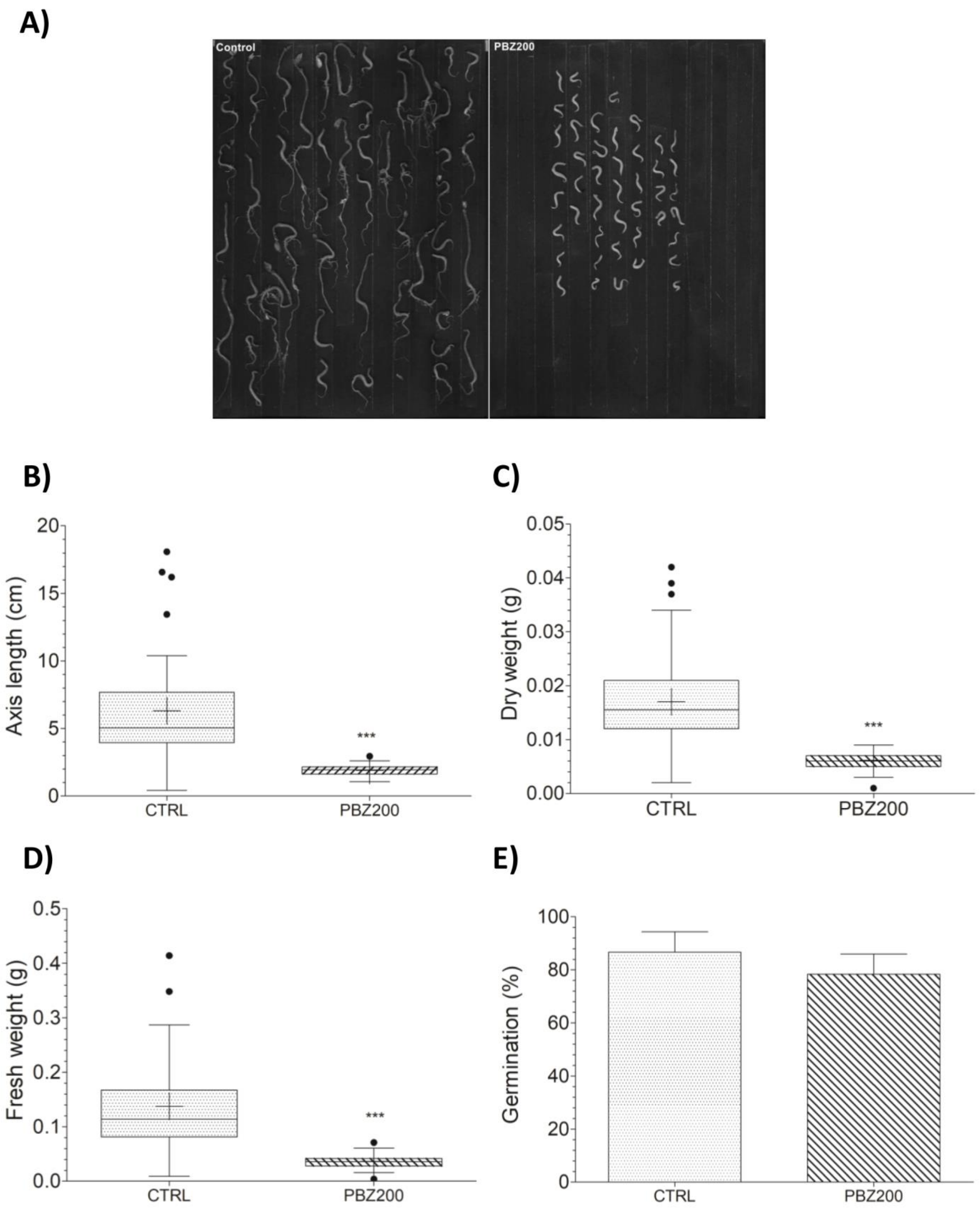


**Supplementary figure S1.** PBZ delays soybean seed germination. Soybean seeds (BRS-284) were allowed to germinate and grow in an incubation chamber under 28°C temperature, and 12/12h photoperiod (dark/light) for 7 days. Seeds were germinated in the presence of 30 ml of sterile water (control) or sterile water with 200 μM paclobutrazol (PBZ). **(A)** Photographs of embryonic axis submitted or not to PBZ treatment. **(B)** Embryonic axis length, **(C)** Dry weight, **(D)** Fresh weight and **(E)** Seed germination. Asterisks above boxplot demonstrate significant difference ($p<0.0001$, Student´s T-test).

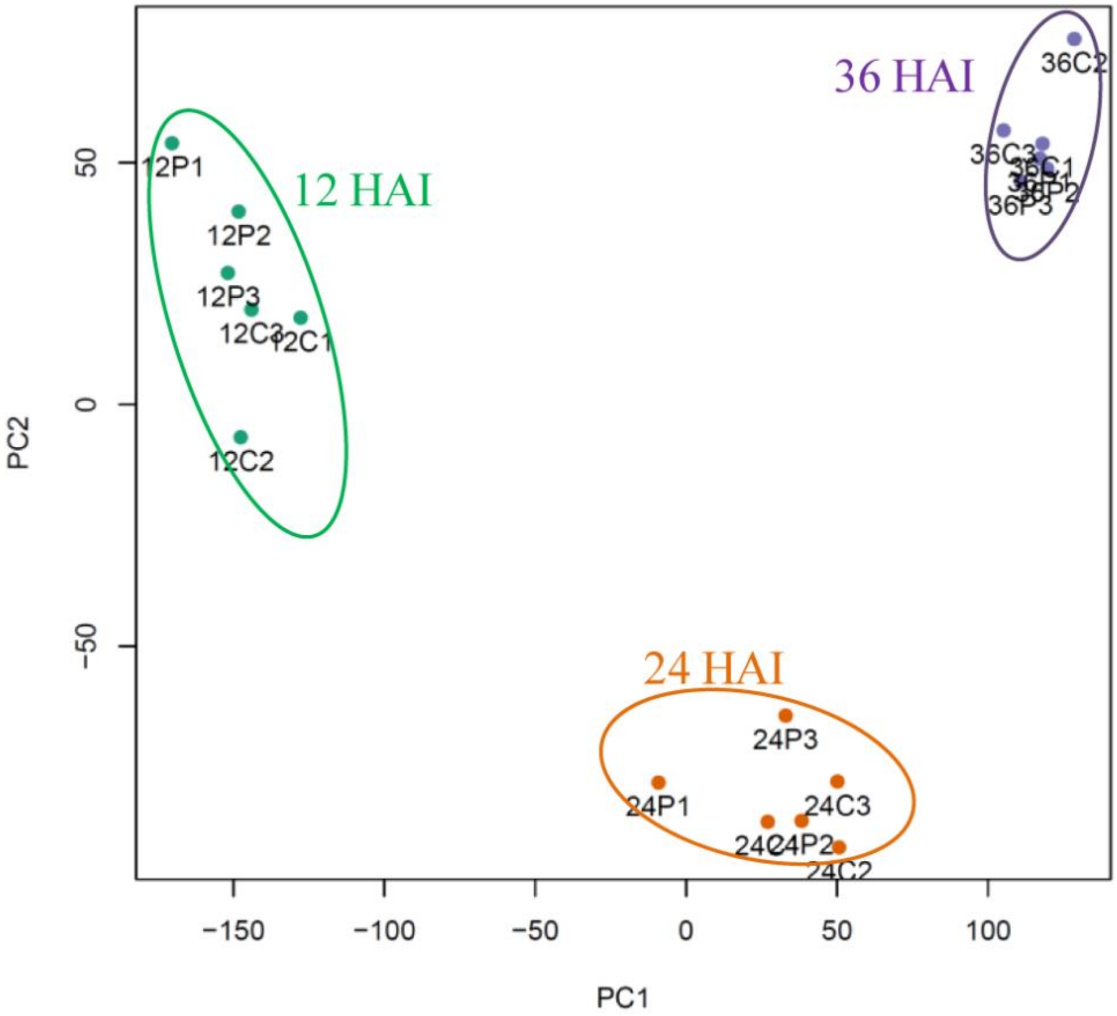


**Supplementary figure S2.** Principal Component Analysis (PCA) of expressed genes under control and PBZ at 12 HAI, 24 HAI and 36 HAI. Three distinct groups can be observed: 12 HAI (green), 24 HAI (orange) and 36 HAI (purple).

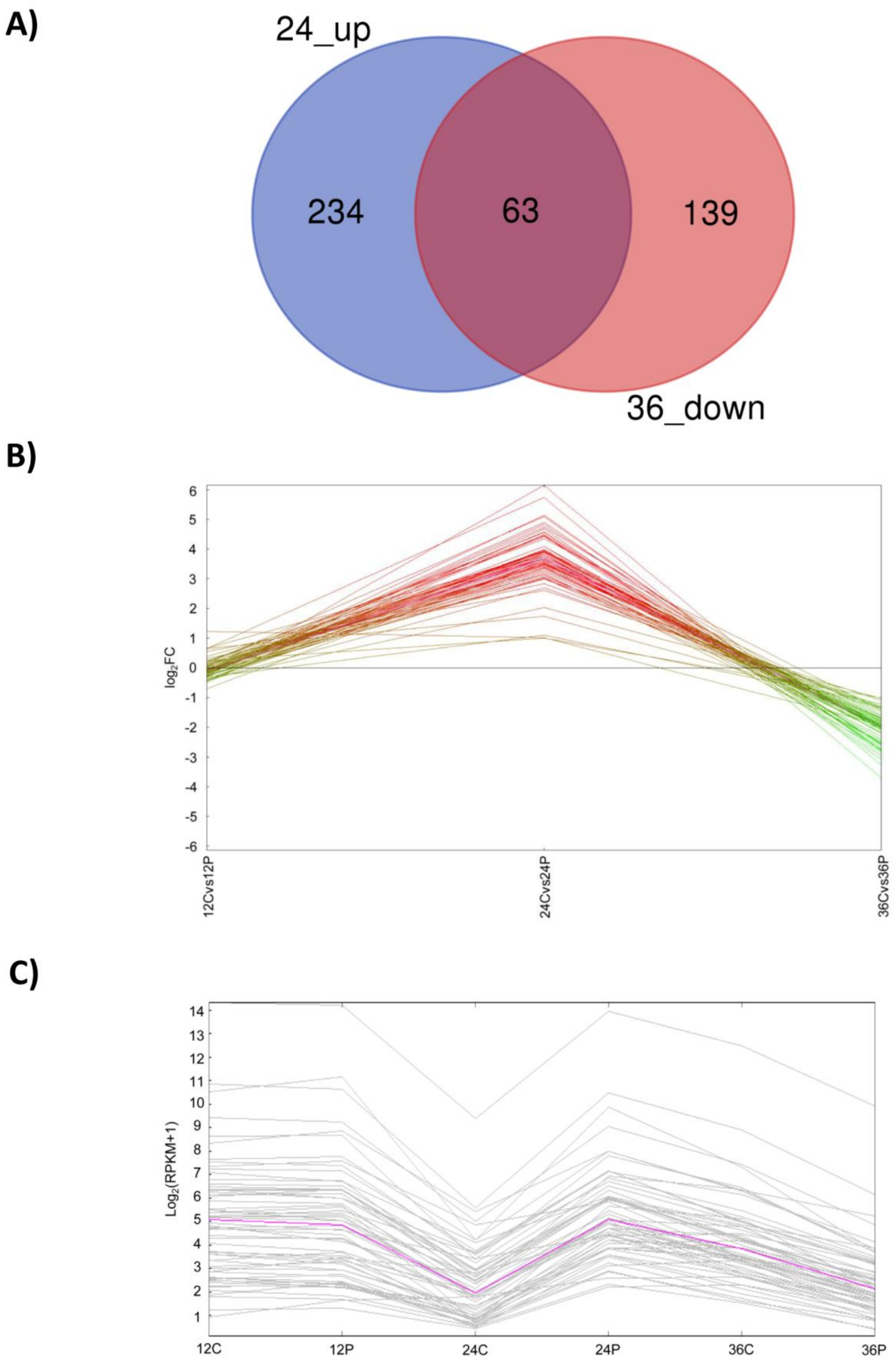


**Supplementary figure S3.** Overlap between 24-up- and 36-down-regulated genes. **(A)** Venn diagram (http://bioinformatics.psb.ugent.be/webtools/Venn/) shows 63 overlapping genes between 24 PBZ up- and 36 down-regulated genes. **(B)** Line plot of $\log_2$(fold change, FC) of the 63 genes at each time point. **(C)** Expression profile of the 63 genes in $\log_2$(RPKM+1) at each time point. MeV was used to generate graphs B and C. The pink line represents the median gene expression.
